# Exploring the microdiversity within marine bacterial taxa: Towards an integrated biogeography in the Southern Ocean

**DOI:** 10.1101/2021.03.03.433356

**Authors:** G Schwob, NI Segovia, CA González-Wevar, L Cabrol, J Orlando, E Poulin

## Abstract

The phylogeography traditionally correlates the genetic relationships among individuals within a macroorganism species, to their spatial distribution. Most microbial phylogeographic studies so far have been restricted to narrow geographical regions, mainly focusing on isolated strains, either obtained by culture or single-strain natural enrichments. However, the laborious culture-based methodology imposes a low number of studied individuals, leading to poor resolution of haplotype frequency estimation, making difficult a realistic evaluation of the genetic structure of natural microbial populations in the environment.

To tackle this limitation, we present a new approach to unravel the phylogeographic patterns of bacteria combining (i) community-wide survey by 16S rRNA gene metabarcoding, (ii) intra-species resolution through the oligotyping method, and (iii) genetic and phylogeographic indices, as well as migration parameters, estimated from populational molecular data as traditionally developed for macroorganisms as models.

As a proof-of-concept, we applied this methodology to the bacterial genus *Spirochaeta*, classically reported as a gut endosymbiont of various invertebrates inhabiting the Southern Ocean (SO), but also described in marine sediment and in open waters. For this purpose, we centered our sampling into three biogeographic provinces of the SO; maritime Antarctica (King George Island), sub-Antarctic Islands (Kerguelen archipelago) and Patagonia in southern South America. Each targeted OTU was chaLRracterized by substantial intrapopulation microdiversity, a significant genetic differentiation and a robust phylogeographic structure among the three distant biogeographic provinces. Patterns of gene flow in *Spirochaeta* populations support the role of the Antarctic Polar Front (APF) as a biogeographic barrier to bacterial dispersal between Antarctic and sub-Antarctic provinces. Conversely, the Antarctic Circumpolar Current (ACC) appears as the main driver of connectivity between geographically distant sub-Antarctic areas such as Patagonia and Kerguelen archipelago, and between Kerguelen archipelago and maritime Antarctica. Additionnally, we found that historical processes (drift and dispersal limitation) together govern up to 86% of the spatial turnover among *Spirochaeta* populations. Overall, our approach represents a substantial first attempt to bridge the gap between microbial and macrobial ecology by unifying the way to study phylogeography. We revealed that strong congruency with macroorganisms patterns at the populational level shaped by the same oceanographic structures and ecological processes.

## Background

Biogeography has traditionally investigated the geographic distribution of macroorganisms in the Eukaryota domain. However, during the last two decades, a growing number of studies has focused on the biogeography of prokaryotic microorganisms, taking advantage of the breakthrough and the constant advances of next-generation sequencing (NGS), which allow extensive surveys of previously inaccessible microbial diversity from a wide range of ecological contexts [1]. Although long debated in the past, it is now accepted that microbes do have biogeographic patterns, repeatedly illustrated by the observation of non-random community assemblages of various prokaryotic microorganisms [2–4]. Contrary to the contemporary driving factors (*i.e.* environmental selection) that have been extensively studied [5–9], the role of historical processes –past ecological and evolutionary events– onto the present-day distribution patterns of microorganisms remains poorly investigated. Initially, the consensus was that the rapid and widespread dispersal of microbes should erase any signal of past historical events [1]. Nevertheless, it is now clear that historical processes, such as the dispersal barriers and geographic distance, might substantially contribute to microbes’ biogeography instead of environmental filtering [5, 10, 11]. For instance, biogeographic regionalization, isolation, and endemism have been reported for microbes, as well as in larger organisms, and evidence a predominant effect of geographic distance over environmental variations [12, 13].

To date, most of the microbial biogeographic patterns have been depicted at the whole community level [4, 14, 15]. Nevertheless, as observed in various empirical studies, a finer taxonomic scale generally allows better detection of geographic patterns [10, 16, 17]. Moreover, the ecological processes driving the biogeographic patterns at the community-level intrinsically result from the accumulation of micro-evolutive processes, *i.e.* mechanisms contributing to the genetic composition and diversity within populations, and how they vary in space and time [10, 18]. The comprehensive description of these micro-evolutive processes requires considering intra-population diversity, as commonly applied in phylogeographic studies of macroorganisms. Hence, microbial assembly processes need to be investigated at a finer taxonomic resolution than usually done by microbial biogeographic surveys and consider the “microdiversity” within groups [17, 18].

The oceans have been considered among the most challenging environments to test hypotheses about microbial biogeography, mainly due to the speculated transport of organisms over large distances by marine currents and the absence of perceivable marine barriers impeding potential dispersal events [19]. However, oceanic fronts separating different water masses have been recently identified as major microbial dispersal barriers [20]. The Southern Ocean (SO) is a vast region representing approximately 20% of the world ocean surface. It surrounds Antarctica, and its northern limit is the Subtropical Front [21]. Two main oceanographic structures shape the SO biogeography; the Antarctic Polar Front (APF) and the Antarctic Circumpolar Current (ACC). The APF is classically considered a harsh north-south obstacle for dispersing marine organisms due to the brutal change in water temperature and salinity [22, 23]. Phylogenetic reconstruction achieved on various vertebrate and invertebrate taxa clearly supports the role of the APF on their respective diversification processes [24–29]. Accordingly, and based on the described distribution of species, biogeographers have traditionally recognized Antarctica and sub-Antarctica as the two main biogeographic provinces of the SO, even if several provinces have been proposed within each of them [30]. Contrarily, outside the APF, the ACC is generally described as the driver of genetic connection across the sub-Antarctic zone due to its clockwise circulation [31–35]. Intraspecific genetic and phylogeographic studies of macroorganisms have demonstrated the ACC’s role in connecting geographically distant sub-Antarctic provinces [32, 36–38]. The marine biota distribution in this region has been synthesized in the Biogeographic Atlas of the SO, providing updated biogeographic information of a wide range of benthic and pelagic taxa from Metazoan, macroalgae, and phytoplankton [39]. Despite being the most abundant and diverse domains on Earth, Bacteria and Archaea were not included in the SO Atlas [40]. Even when marine microbial communities have been previously characterized in the region, their geographic distribution’s underlying drivers remain poorly understood. Studies conducted at the whole community-level support (i) the role of ACC as a likely efficient mechanism of circumpolar microbial transport and dispersal [3, 41] and (ii) the role of APF as the main dispersal barrier separating Antarctic and sub-Antarctic microbial assemblages [3, 20, 42]. However, the observed biogeographic patterns’ underlying evolutionary processes have not been elucidated and would intrinsically rely on bacterial populations’ microdiversity.

Targeting the intraspecific diversity using NGS data requires specific computational methods to discriminate the stochastic noise caused by random sequencing errors from those associated with biologically significant diversity [10, 17, 43]. For this purpose, an algorithm called “Minimum Entropy Decomposition” (MED) relying on the oligotyping method has been proposed by Eren *et al*. [44]. This algorithm allows to identify true sequence variants (*i.e.* oligotypes) within the “Operational Taxonomic Units” (OTUs), classically defined at 97% identity of the bacterial 16S rRNA gene. This approach has already been successfully used to unravel fine-grained biogeographic patterns of bacterial microdiversity in Arctic sediments, such as variations in oligotype distribution according to spatial and environmental parameters [45]. Moreover, focusing on the sulfate-reducing genus *Desulfotomaculum* in Arctic marine sediments, Hanson *et al.* [11] showed clear biogeographic patterns –attributed to historical factors associated with past environments– were only evident at the microdiversity level achieved with the oligotyping method. However, the microevolutionary processes causing the microdiversity were not assayed, and the study did not encompass large-scale distribution among different biogeographic provinces, as it was restricted to the west coast of Spitsbergen, Svalbard in the Arctic Ocean.

In the present proof-of-concept study, we aim to bring new insights on the evolutionary processes driving microbial biogeography across different provinces of the SO by combining (i) community-wide surveying provided by the high-throughput sequencing of the 16S rRNA gene, (ii) intra-species resolution obtained through the oligotyping method implemented in the MED pipeline, and (iii) phylogeographic analysis as traditionally developed for macroorganisms as models. Considering the SO as an outstanding idoneous frame, we investigated the geographic distribution of genetic diversity of marine bacterial taxa across three main biogeographic provinces: maritime Antarctica (King George Island, South Shetland Islands, West Antarctic Peninsula), sub-Antarctic Islands of the Indian Ocean (Kerguelen archipelago) and southern South America (Patagonia), encompassing sites separated by the APF, and others connected through the ACC.

As the contribution of geography to biological diversity patterns (*i.e.* dispersal limitation) is stronger on habitat-specialists (*i.e.* taxa found in habitat with high selective strength) [46, 47], and emphasized within homogeneous habitats distributed across large spatial scales [10, 48, 49], we focused our study on the bacterial community associated to a specific habitat: the gut of *Abatus* irregular sea urchins. The *Abatus* genus is distributed across the SO and gathers various sibling species homologous in ecology and habitat, such as *Abatus cavernosus* in southern South America, *Abatus cordatus* in Kerguelen Islands, and *Abatus agassizii* in maritime Antarctica [50–53]. Since these species lack specialized respiratory structure, they are restricted to the well-oxygenated coarse sediments found at shallow depth (typically 1 to 3 meters depth) in sheltered bays, protected from the swell [51]. Within the *Abatus* hosts, we focused on a specific micro-environment –the gut tissue– previously described to act as a selective filter of the external sediment microbiota, as illustrated by the reduction of bacterial diversity at both taxonomic and functional levels [54]. Working on the gut community with supposedly more limited dispersal capacity, through a high sequencing depth, is expected to (i) provide robust coverage of the bacterial diversity, (ii) minimize the relevance of environmental filtering between provinces, (iii) emphasize the contribution of geographic and oceanographic factors, and therefore (iv) enhance the detection of phylogeographic signals across the SO [10]. As a model taxon to explore bacterial phylogeography in the SO, we chose the *Spirochaeta* genus (phylum *Spirochaetes*). *Spirochaeta* bacteria are recognized as the most prevalent and abundant genus in the *Abatus* gut tissue [54]. Moreover, spirochaetes are classically found in marine benthic sediments [55, 56] and, to a lesser extent, in the water column (Ocean Barcode Atlas; http://oba.mio.osupytheas.fr/ocean-atlas/). Thus, due to its ease of detection and ubiquity across biogeographic provinces, *Spirochaeta* represents an illustrative model to validate our methodology and explore marine bacteria’s spatial genetic patterns, from genus to population level. We hypothesized that the strong biogeographic barrier between South America and maritime Antarctica classically observed in the literature for macroorganisms (*i.e.* vicariance process) also affects the fine-scale genetic structure and the phylogeographic patterns within *Spirochaeta* OTUs. Besides, the ACC-mediated connectivity among sub-Antarctic provinces should be reflected by a greater genetic homogeneity of *Spirochaeta* populations between South American sites and the Kerguelen Islands, rather than with maritime Antarctica. Alternatively, the potential high dispersal capacity of *Spirochaeta* taxa may result in the absence of genetic and phylogeographic structure across the SO.

## Methods

### Sampling collection, DNA extraction, and 16S rRNA gene library preparation

Adult *Abatus* individuals were sampled from four localities across the SO, including two sites in Patagonia, southern South America (Possession Bay, PAT1 and Puerto Deseado, PAT2), one site in Kerguelen Islands (Port-aux-Français, KER), and one site in the West Antarctic Peninsula (King George Island, KGI) (Figure 1, Table 1).

**Figure 1.**
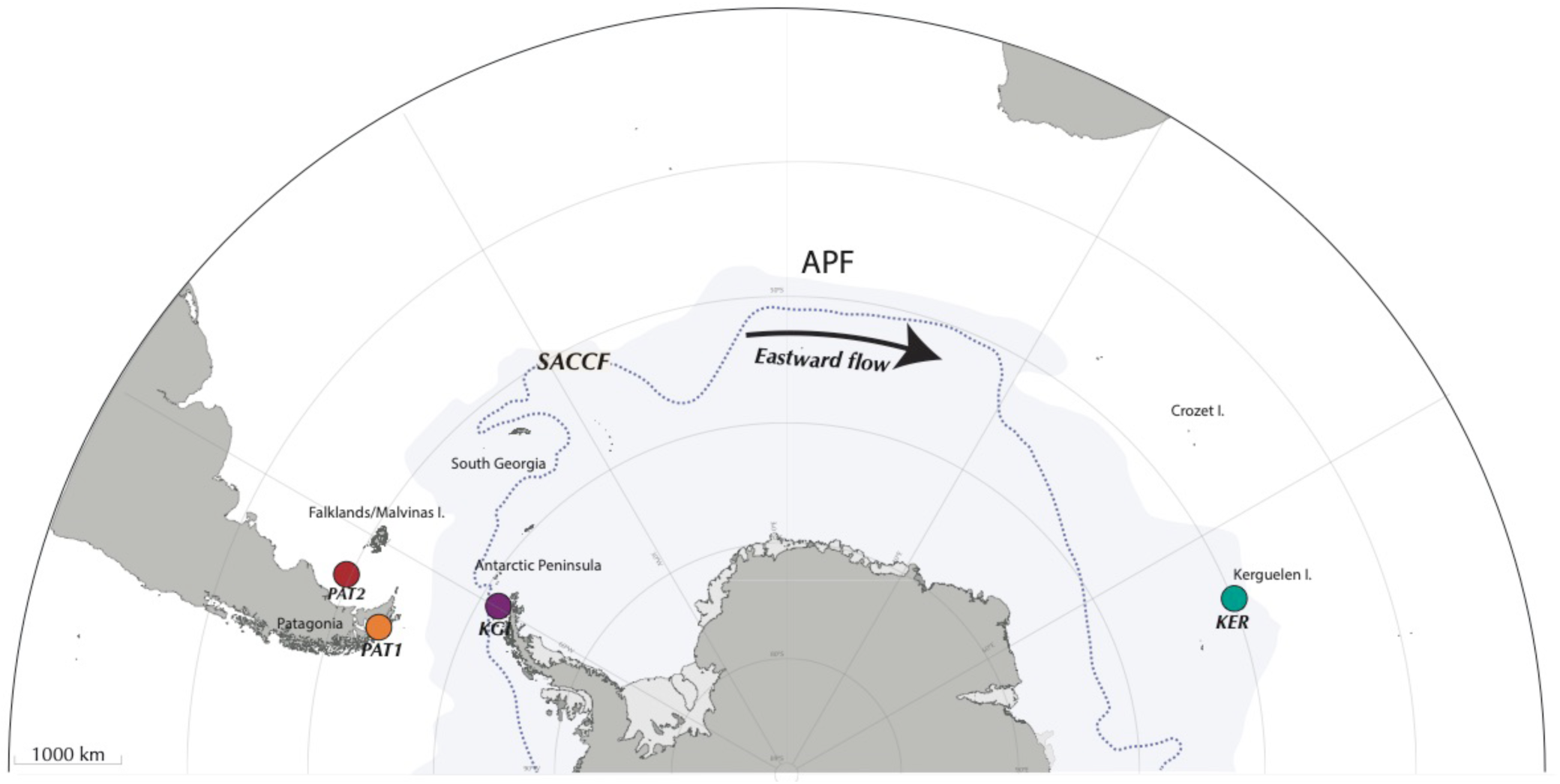
Sampling localities across the Southern Ocean, encompassing Possession Bay and Puerto Deseado in Patagonia (PAT1 and PAT2, respectively), King George Island in Maritime Antarctica (KGI), and Port-aux-Français in Kerguelen Islands (KER). The Antarctic Polar Front (APF) and the Southern Antarctic Circumpolar Front (SACCF) are represented.

**Table 1.** Experimental design and sequencing data

| Locality | Province | GPS coordinates | Date | Designation | Sample types | N | Nseq. (Relat. Abund.) |
| --- | --- | --- | --- | --- | --- | --- | --- |
| King George Island | Maritime Antarctica | 62°12'55.3"S | 01-2019 | KGI | External sediments | 8 | 255786 (10%) |
|  |  | 58°56'43.8"W |  |  | Gut tissue | 31 | 563383 (22%) |
| Bahía Posesión | Patagonia | 52°19'52.97"S | 07-2019 | PAT1 | External sediments | 6 | 271828 (11%) |
|  |  | 69°29'10.50"W |  |  | Gut tissue | 15 | 447892 (18%) |
| Puerto Deseado | Patagonia | 47°45'07.0"S | 12-2016 | PAT2 | External sediments | NA | NA |
|  |  | 65°52'04.0"W |  |  | Gut tissue | 10 | 470087 (18%) |
| Port-aux-Français | Kerguelen Island | 49°21'13.32"S | 11-2017 | KER | External sediments | 5 | 92564 (4%) |
|  |  | 70°13'8.759"E |  |  | Gut tissue | 14 | 440498 (17%) |

N: number of samples, Nseq.: total number of cleaned sequences obtained, Relat. Abund.: (relative abundance in the global dataset).

Marine surface sediments (0 – 5 cm, referred here as “external sediment”) were also sampled in each *Abatus* population’s immediate vicinity as the ingested food source of the sea urchins. Due to logistic constraints, it was not possible to collect external sediment in the PAT2 site. All individuals were dissected under sterile conditions to collect the whole digestive tract minus the caecum (identified as “gut tissue”) following Schwob *et al.* [54]. Gut tissue samples were gently rinsed with nuclease-free sterile water to remove the content (*i.e.* in sediment) and were then individually homogenized using mortar and pestle under a laminar-flow cabinet. Genomic DNA was extracted from the totality of the homogenized gut tissue samples using the DNeasy PowerSoil^®^ Kit (Qiagen, Hilden, Germany) following the manufacturer’s recommendations.

A metabarcoding approach was used to assess the bacterial community composition in the external sediment and *Abatus* gut tissue samples. Briefly, extracted genomic DNA was used as the template for PCR amplification using the primers 515F 5’-GTGYCAGCMGCCGCGGTA-3’ and 926R 5’-CCCCGYCAATTCMTTTRAGT-3’ [57]. The PCR conditions and 16S rRNA gene library preparation were the same as described in Schwob *et al.* [54].

### Metabarcoding data processing

External sediment and gut tissue amplicons were sequenced using the paired-end sequencing technology (2 x 250 bp) on the Illumina MiSeq platform at the UWBC DNA Sequencing Facility (University of Wisconsin-Madison, USA). Reads of 16S rRNA were processed using the open-source software MOTHUR v1.44.0 following Schwob *et al.* [54]. Once the raw reads were processed into Operational Taxonomic Units (OTUs) at 97% identity threshold, we applied a filter of relative abundance at > 0.0001% as recommended by Bokulich *et al.* [58]. Following this, a taxonomic classification was performed with the *classify.otu* function and the SILVA database v138 implemented in MOTHUR. An OTU table of *Spirochaeta* was edited (*i.e.* all OTUs assigned to the genus *Spirochaeta*), and converted into a presence/absence matrix. Bray-Curtis and Unweighted Unifrac distances were calculated from the OTU presence/absence matrix and used to perform a Non-Metric Multidimensional Scaling (NMDS) with the *metaMDS* function of the ADE4 package [59]. The NMDS was plotted through a scatter diagram using the *s.class* function implemented in ADE4. The locality’s contribution to the *Spirochaeta* OTUs composition in gut tissue samples was tested with permutational multivariate analysis of variance (permanova), using *adonis* and *pairwise.adonis* functions implemented in *vegan* and *pairwiseAdonis* R packages, respectively [60, 61].

A subset of the three most abundant *Spirochaeta* OTUs present in the four localities was retained for further analysis (Table S2). All the sequences assigned to the selected *Spirochaeta* OTUs were retrieved using the *bin.seqs* command in MOTHUR. Finally, the resulting fasta files were processed independently through the Minimum Entropy Decomposition pipeline following Eren *et al.* [44].

### Minimum Entropy Decomposition

Minimum Entropy Decomposition (MED) pipeline was used to identify nucleotidic polymorphism at fine-scale resolution (> 3% identity) within the 16S rRNA gene sequences from *Spirochaeta* OTUs. Briefly, MED employs the Shannon entropy algorithm to discriminate biologically meaningful variations of closely related sequences from the stochastic noise caused by random sequencing errors, focusing on informative-rich variable nucleotide positions [44, 62]. The resulting taxonomic units will be referred to as *Spirochaeta* oligotypes. Unsupervised oligotyping was carried out individually on *Spirochaeta* OTUs using the default dynamically computed threshold from which entropy is considered as zero (-m). Additionally, each identified oligotype had to have a default minimum relative abundance of 2% in the OTU sequences dataset [44]. Accumulation curves of oligotypes’ richness were computed for each *Spirochaeta* OTU at a 95% confidence interval using the package INEXT [63] in R v3.6.0 [64]. Pie charts of the relative site contributions in the total abundance of the *Spirochaeta* OTUs oligotypes were performed with the *pie* function in the package GRAPHICS in R v3.6.0.

### Genetic diversity and structure of *Spirochaeta* populations

The number of oligotypes (*k*), the oligotype diversity (*H*), the number of discriminant sites (*S*) and the pairwise difference between sequences (*Π*) were estimated individually for each *Spirochaeta* OTU using the packages PEGAS [65] and APE v5.3.0 [66] in R v3.6.0. For comparative purposes among sites with unequal sample sizes, a composite bootstrapping script was written to rarefy the sequence datasets to the minimum number of sequences per site and repeat the rarefaction with 1,000 re-samplings. Confidence intervals at 95% of genetic diversity indices were then calculated using these iteration values. The genetic differentiation (F_ST_ and ɸ_ST_) among *Spirochaeta* populations was analyzed using the software ARLEQUIN v3.5.2 [67] with 1,000 permutations and a significance threshold at 0.05. Phylogeographic differentiation was also estimated with the nearest-neighbor statistic Snn [68], and the significance of Snn estimates was tested with a permutation test through DNASP v5.10.01 [69]. The reconstruction of oligotype networks was performed using the Median Joining method with the software Populational Analysis with Reticulate Trees v1.7.0 in PopART [70]. Oligotype abundances were normalized by the number of sequences per locality for a given OTU to improve the networks’ readability.

### Quantification of selection, dispersal, and drift

The relative contribution of stochastic (*i.e.* dispersal, drift) and deterministic (*i.e.* selection) processes, also referred to as historical and contemporary processes respectively, on *Spirochaeta* oligotype assembly was measured for the selected OTU by following the analytical framework described in Stegen *et al.* [71] and illustrated by Feng *et al.* [72]. In a nutshell, the approach relies on the comparison of the phylogenetic turnover between communities across samples (βMNTD; β mean nearest-taxon distance) to a null distribution of βMNTD, and denoted as the β-nearest taxon index (βNTI). The phylogenetic tree required for the βMNTD/βNTI calculations was generated using PhyML v3.0 [73], and the oligotype sequences of *Spirochaeta* previously aligned with MUSCLE [74]. βNTI values indicate that taxa between two communities are more (*i.e.* βNTI < -2) or less (*i.e.* βNTI > +2) phylogenetically related than expected by chance, thus suggesting that communities experience homogenizing or variable selection, respectively [75]. βNTI values ranging from -2 to +2 indicate a limited selection effect and point to dispersal limitation and ecological drift out as possible community composition drivers. To further disentangle the respective effect of these two alternatives processes, we calculated the pairwise Bray Curtis-based Raup-Crick dissimilarity index (RC_Bray_) between sites [76], weighted by oligotype abundance [71]. For this, we used an optimized version of the initial script of Stegen *et al.* [71], developed by Richter-Heitmann [77], and available via GitHub (https://github.com/FranzKrah/raup_crick). RC_Bray_ values < - 0.95 and > + 0.95 correspond to communities that have –respectively– more or fewer taxa in common than expected by chance, and therefore indicate that community turnover is driven by homogenising dispersal (RC_Bray_ < - 0.95) or dispersal limitation plus drift (RC_Bray_ > + 0.95). On the contrary, RC_Bray_ values > - 0.95 and < + 0.95 are indicative of ecological drift [78]. Both βNTI and RC_Bray_ null models included 999 randomizations [71].

### Testing for isolation by distance (IBD) and environment (IBE)

To disentangle the relative effect of geographic distance and abiotic environmental differences on the *Spirochaeta* oligotype composition between samples, we used the distance-based multiple matrix regression with randomization (MMRR) approach [79]. We extracted a set of 9 environmental variables for each of our sampling site from the Bio-ORACLE database [80], including pH, the means of nitrate, silicate, and phosphate concentrations, and the means at the mean depth of seawater salinity, dissolved oxygen concentration, seawater temperature, seawater temperature range and chlorophyll concentration. All environmental variables were standardized ((*x_i_ − x̄)/sd(x)*), and were then analyzed through Principal Components Analysis (PCA). As a high percentage of the variation among localities was explained by the first component (PC1, >91%), we transformed the scores of PC1 into Euclidean distance using the *vegdist* function in the *vegan* package in R to use it as the environmental distance matrix further. The longitude and latitude coordinates were converted into kilometers using the *earth.dist* function implemented in the FOSSIL package in R [81]. The geographic distances were transformed using the Hellinger method through the *decostand* function of the *vegan* package in R. The dissimilarity matrix of *Spirochaeta* oligotype composition among samples was also created from Bray-Curtis distances using the *vegdist* function of the R package *vegan*. Finally, to evaluate the relative weight of environmental and geographic distance matrices, an MMRR was performed using the R package PopGenReport [82], and the correlation coefficients and their significance were estimated based on 10,000 random permutations.

### Connectivity among *Spirochaeta* populations

The amount and direction of gene flow among *Spirochaeta* populations were estimated using the coalescent-based program LAMARC v2.1.10 [83]. A total of ten runs was performed for each *Spirochaeta* OTU, consisting of likelihood searches of 20 initial and two final chains, with a minimum of 500 and 10,000 recorded trees, respectively, and sampling every 20 generations after a burn-in of 1,000 genealogies. The effective number of migrants per generation (Nm) among *Spirochaeta* populations was calculated by multiplying the Maximum Likelihood Estimates (MLE) of the mutation parameter (θ) by the migration parameter (M), both estimated through the LAMARC program. We present the mean and standard deviation of the estimated Nm values obtained from the ten runs for each *Spirochaeta* OTU.

## Results

### Sequencing performance and OTUbased analysis

A total of 4,184,226 raw reads was generated from the 91 samples of external sediments and gut tissues. Once processed, 2,542,038 cleaned sequences distributed into 727 OTUs were obtained from the external sediment and gut tissue samples (details provided in Table 1). Out of this condensed dataset, 425,613 sequences associated with the *Spirochaeta* genus were retrieved, representing a total of 10 OTUs.

Both Bray-Curtis and Unweighted Unifrac distance methods did not reveal any difference in *Spirochaeta* OTU composition between the Patagonian sites (PAT1 and PAT2) (Figure 2A and 2B, Additional file 1).

**Figure 2.**
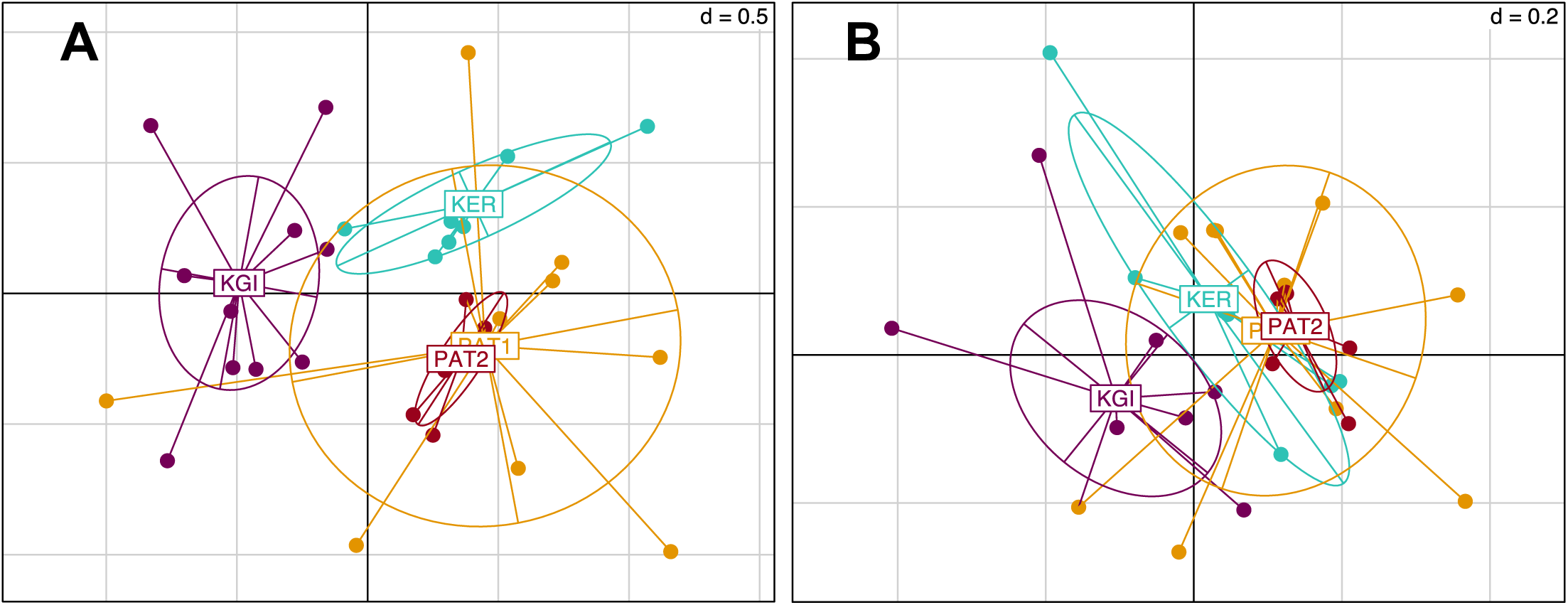
NMDS scatter diagram of the *Spirochaeta* OTUs composition in gut tissue samples across the localities. **(A)** Presence/absence matrix converted in Bray-Curtis distances, **(B)** Unweighted Unifrac distance. Colors are assigned to the locality.

Conversely, Kerguelen Islands (KER) and martime Antarctic (KGI) sites were significantly different from each other in terms of *Spirochaeta* OTU composition and with the Patagonian ones (Additional file 1).

The relative abundance analyses among the 10 *Spirochaeta* OTUs (Figure 3) showed four of them were shared among all the Southern Ocean’s sampled provinces. Three OTUs (OTU6, OTU7, and OTU532) were more abundant in maritime Antarctica (KGI), three (OTU23, OTU42, and OTU221) were predominantly found in the Patagonian locality PAT2, three (OTU40, OTU278, and OTU561) were more abundant in the Patagonian locality PAT1, and a single one (OTU349) was predominant in Kerguelen Islands (Figure 3).

**Figure 3.**
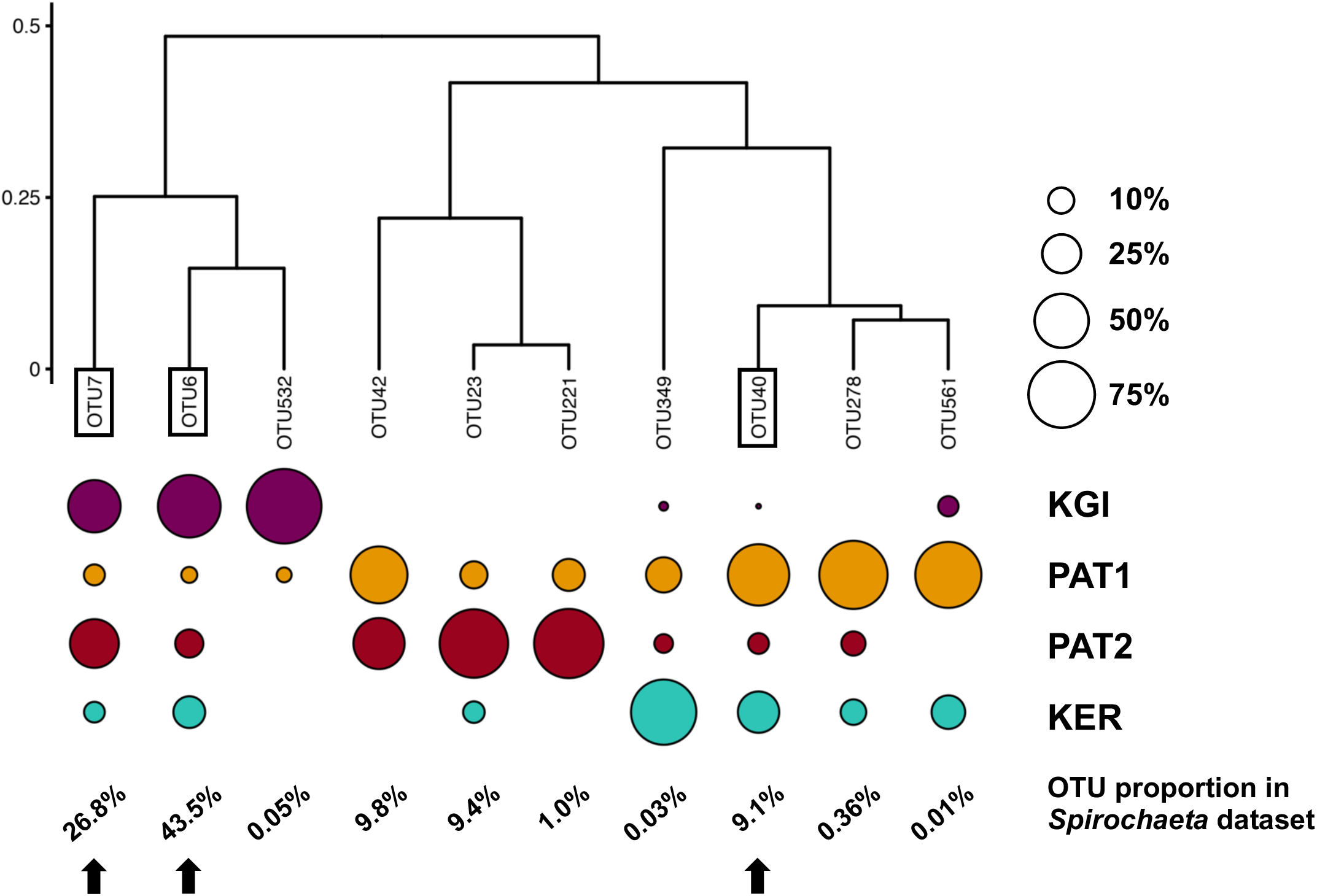
Clustering of *Spirochaeta* OTUs based on their relative abundance in each site. Clustering of Bray-Curtis distances matrix using the complete linkage method. The size of the circles indicates the repartition of a given OTU sequences among the 4 localities. The percentages indicate the OTUs’ proportions in the *Spirochaeta* dataset. Framed labels and black arrows indicate the selected OTUs that were selected to be process through the Minimum Decomposition Entropy pipeline (MED).

To test the genetic and phylogeographic structure of *Spirochaeta* at the broadest geographic scale available through our dataset, we selected the three most abundant *Spirochaeta* OTUs which were detected within the four localities of the dataset (*i.e.* OTU6, OTU7, and OTU40). These three selected co-distributed OTUs are good targets to constitute a metapopulation, which is a meaningful ecological unit of distinct local populations separated by gaps in habitats and interconnected to some extent via dispersal events of individuals [84]. The relative abundance in sample types, the closest sequence retrieved from Blast analysis, and the distribution of these OTUs among the localities are provided in the Additionnal files 2 and 3.

### Microdiversity within *Spirochaeta* OTUs

A total of 48, 96, and 48 oligotypes were defined for OTU6, OTU7, and OTU40, respectively (Additionnal file 4). Accumulation curves of OTU6, OTU7, and OTU40 oligotypes reached saturation in almost all localities indicating that the overall majority of *Spirochaeta* microdiversity has been found within the analyzed samples (Additionnal file 5). Diversity indices measured as *N*, *k*, *S*, *h,* and Π for each OTU in each locality are provided in Table 2.

**Table 2.** Summary of oligotypes number and genetic indices per OTU and per site for the three most abundant *Spirochaeta* OTUs found in all sampling localities.

| OTU | Site | $N$ | $k$ | $S$ | $H$ | $\Pi$ |
| --- | --- | --- | --- | --- | --- | --- |
| OTU6 | KGI | 125,112 | 26 ± 0 | 24 ± 0 | 0.4879 ± 0.0004 | 0.7919 ± 0.0012 |
|  | PAT1 | 6,452 | 18 ± 0 | 8 ± 0 | 0.6701 ± 0.0000 | 1.5772 ± 0.0000 |
|  | PAT2 | 22,545 | 22 ± 0 | 12 ± 0 | 0.6574 ± 0.0002 | 1.5426 ± 0.0006 |
|  | KER | 29,448 | 33 ± 1 | 29 ± 0 | 0.5677 ± 0.0003 | 0.9687 ± 0.0011 |
| OTU7 | KGI | 53,645 | 31 ± 0 | 32 ± 0 | 0.5555 ± 0.0003 | 1.4443 ± 0.0021 |
|  | PAT1 | 7,249 | 69 ± 0 | 37 ± 0 | 0.8036 ± 0.0000 | 2.0247 ± 0.0003 |
|  | PAT2 | 44,509 | 60 ± 0 | 33 ± 0 | 0.7958 ± 0.0002 | 1.7336 ± 0.0011 |
|  | KER | 7,021 | 43 ± 0 | 37 ± 0 | 0.6306 ± 0.0000 | 1.0261 ± 0.0000 |
| OTU40 | KGI | 47 | 4 ± 0 | 4 ± 0 | 0.6984 ± 0.0000 | 1.6606 ± 0.0000 |
|  | PAT1 | 24,612 | 32 ± 0 | 14 ± 0 | 0.3741 ± 0.0007 | 0.4922 ± 0.0011 |
|  | PAT2 | 2,423* | 34 ± 0 | 11 ± 0 | 0.3816 ± 0.0000 | 0.4903 ± 0.0000 |
|  | KER | 11,017 | 12 ± 0 | 8 ± 0 | 0.0681 ± 0.0004 | 0.0913 ± 0.0006 |

$N$ : number of sequences, $k$ : number of oligotypes, $S$ : number of polymorphic sites, $H$ : genetic diversity, $\Pi$ : mean number of pairwise diversity. The mean and standard deviation were calculated from a total of 1,000 bootstraps, performed by randomly subsampling per site a number of sequences equal to the minimum number of sequences obtained among sites for a given OTU. \* In the case of OTU40, the number of sequences in the PAT2 site was used to perform the resampling.

The genetic diversity (*H*) ranged from 0.07 (OTU40 in KER) to 0.80 (OTU7 in PAT1) across localities (Table 2). Patagonian sites exhibited higher oligotype and nucleotide diversity for OTU6 and OTU7 than maritime Antarctica and Kerguelen Islands localities. In contrast, the genetic diversity of the OTU40 oligotypes was higher in the maritime Antarctic site and lower for the Kerguelen population.

### Populations differentiation and phylogeographic structure of *Spirochaeta* oligotypes

Independently of the OTU considered, the genetic (F_ST_) and phylogeographic (ɸ_ST_) structures between the two closest localities (*i.e.* Patagonian sites PAT1 and PAT2) were extremely to moderately low. In the case of the OTU40, the genetic diversity and frequencies of *Spirochaeta* oligotypes were fully homogenous between PAT1 and PAT2, as indicated by the non-significant values of F_ST_ and ɸ_ST_ comparisons (Table 3).

**Table 3.** Genetic (F_ST_) and phylogeographic structure (ɸ_ST_) of the *Spirochaeta* populations among localities.

| OTU | Index | Locality | KGI | PAT1 | PAT2 | KER |
| --- | --- | --- | --- | --- | --- | --- |
| OTU6 | FST | KGI | - | 0 | 0 | 0 |
|  |  | PAT1 | 0.4428 | - | 0 | 0 |
|  |  | PAT2 | 0.4417 | 0.0026 | - | 0 |
|  |  | KER | 0.0654 | 0.3856 | 0.3829 | - |
| | $\phi_{ST}$ | KGI | - | 0 | 0 | 0 |
|  |  | PAT1 | 0.5371 | - | 0 | 0 |
|  |  | PAT2 | 0.5360 | 0.0027 | - | 0 |
|  |  | KER | 0.1041 | 0.5063 | 0.4933 | - |
| OTU7 | FST | KGI | - | 0 | 0 | 0 |
|  |  | PAT1 | 0.3436 | - | 0 | 0 |
|  |  | PAT2 | 0.3254 | 0.0726 | - | 0 |
|  |  | KER | 0.3902 | 0.0568 | 0.1734 | - |
| | $\phi_{ST}$ | KGI | - | 0 | 0 | 0 |
|  |  | PAT1 | 0.4393 | - | 0 | 0 |
|  |  | PAT2 | 0.5361 | 0.1341 | - | 0 |
|  |  | KER | 0.3967 | 0.1312 | 0.3429 | - |
| OTU40 | FST | KGI | - | 0 | 0 | 0 |
|  |  | PAT1 | 0.5549 | - | 0.4505 | 0 |
|  |  | PAT2 | 0.5445 | <0.0001 | - | 0 |
|  |  | KER | 0.8491 | 0.7358 | 0.8614 | - |
| | $\phi_{ST}$ | KGI | - | 0 | 0 | 0 |
|  |  | PAT1 | 0.8326 | - | 0.2793 | 0 |
|  |  | PAT2 | 0.8278 | <0.0001 | - | 0 |
|  |  | KER | 0.9354 | 0.7398 | 0.8652 | - |

Structure values are beneath each diagonal and $p$ -values are above them. $p$ -values were obtained through 1,000 permutations and the significance level was set < 0.05. $p$ -values of 0 indicate value < 0.00001.

Contrarily, higher values of F_ST_ and ɸ_ST_ comparisons were recorded among the three provinces considered in this study (Patagonia, PAT1/PAT2; maritime Antarctica, KGI; Kerguelen Islands, KER) (Table 3). Consistently, the distribution of the *Spirochaeta* oligotypes was geographically discontinuous across the localities, with various province-specific oligotypes (Additionnal file 4). Two exceptions were observed in the case of maritime Antarctica and Kerguelen Islands (KGI and KER) comparisons for both OTU6 and OTU7, with relatively lower values of F_ST_ and ɸ_ST_ (Table 3). The Snn test for phylogeographic structure among all sites was significant with statistic values ≥ 0.5 (*i.e.* OTU6; Snn = 0.50, *p*-value < 0.0001, OTU7; Snn = 0.57, *p-value* < 0.0001, OTU40; Snn = 0.50, *p*-value < 0.0001). All in all, these results indicate the existence of both genetically and geographically differentiated *Spirochaeta* populations across the three biogeographic provinces sampled.

Within the 48 oligotypes identified in the OTU6, 11 (∼23%) were private to one of the three provinces (Patagonia, maritime Antarctica, or the Kerguelen Islands), and more than half were exclusive to Kerguelen Islands (Additionnal file 4). Maritime Antarctic and Kerguelen Islands (KGI and KER) shared 27 (∼66%) of their oligotypes. The Patagonian localities (PAT1/PAT2) shared 18 of the 23 total oligotypes (∼78%) observed in this province (Figure 4, Table S3). A total of five (∼10%) oligotypes were broadly distributed across all localities. Oligotype network of OTU6 showed short genealogies and the presence of at least five dominant oligotypes. The dominant oligotype in Patagonia (PAT1/PAT2), and the dominant oligotype in maritime Antarctica and Kerguelen Islands (KGI/KER), were separated by a single substitution (Figure 4).

**Figure 4.**
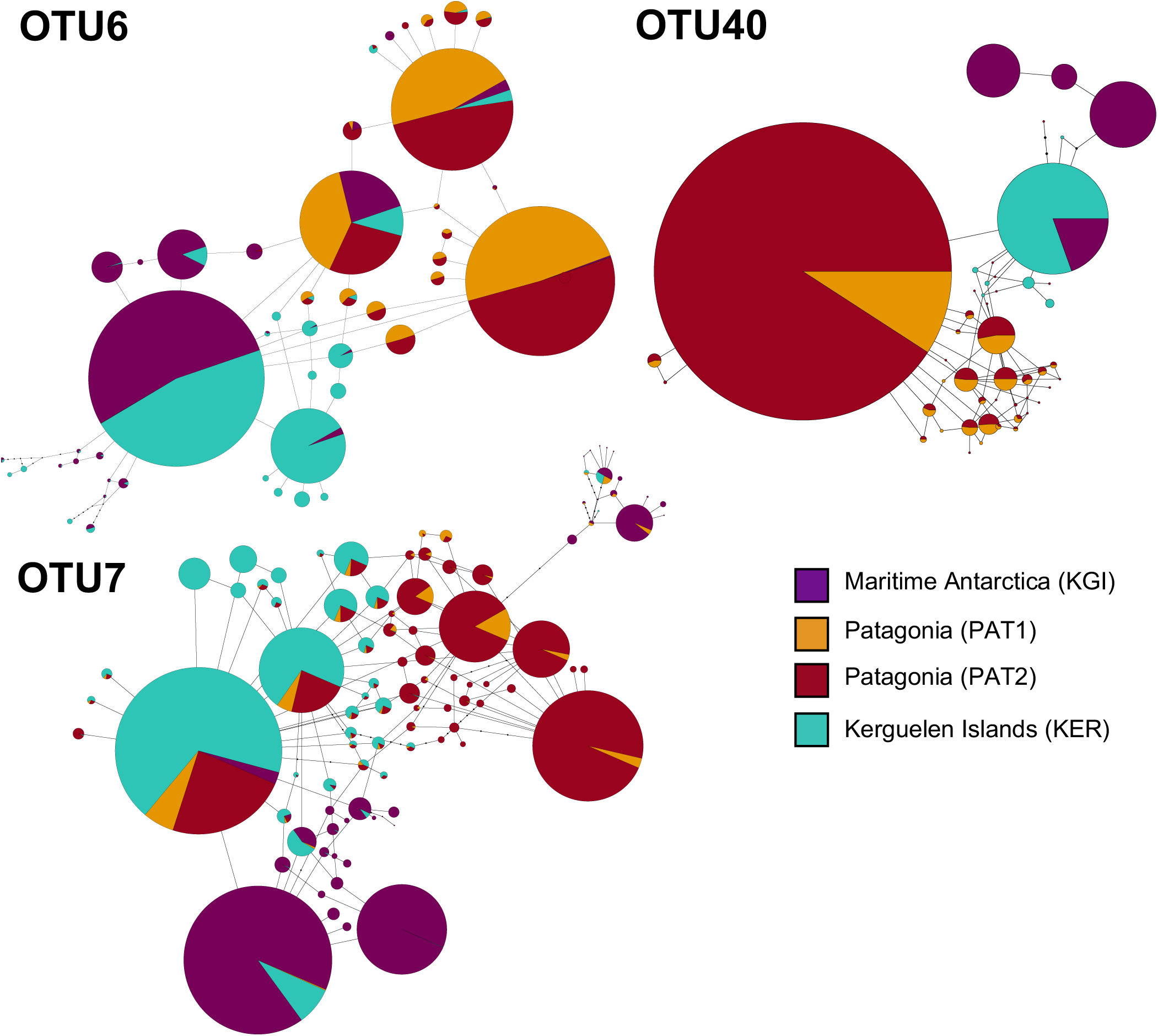
Median-joining oligotype networks of the three selected OTUs of *Spirochaeta*. Each circle represents a distinct oligotype. Colors indicate the locality of provenance. Circle size is scaled on the oligotype frequency normalized by the number of sequences in each locality, within the corresponding OTU dataset. Raw abundances are provided in Table S3.

In the case of OTU7, the percentage of private oligotypes was almost the same as the OTU6, with 21 (∼22%) oligotypes exclusive to one of the provinces (Additionnal file 4). The dominant oligotype was different between maritime Antarctic and Kerguelen Islands localities (KGI and KER) (Figure 4). While most oligotypes from Kerguelen Islands (KER) were detected in at least one of the Patagonian sites (PAT2 or PAT1) (∼94%), fewer oligotypes from maritime Antarctica (KGI) were observed in Kerguelen Islands (∼68%) and even fewer in the Patagonian localities (∼62%) (Figure 4).

For OTU40, we recorded a predominant group of oligotypes specific to the Patagonian sites representing 65% of the oligotypes identified within the OTU40 (Additionnal file 4, Figure 4). A clear separation was observed in the oligotype network between the KGI/KER and PAT1/PAT2 localities (Figure 4), with only 4 shared oligotypes (∼8%) (Table S3). In maritime Antarctica (KGI), three of the four oligotypes were private, whereas the dominant one from Kerguelen Islands (KER) was shared with maritime Antarctica (KGI) (∼8%) (Figure 4, Additionnal file 4).

### Gene flow under a migration–drift equilibrium model

All the analyzed OTUs showed high genetic similarities between the analyzed Patagonian populations (PAT1 and PAT2). Gene flow analyses identified a bidirectional pattern from PAT2 to PAT1 (effective number of migrants per generation, Nm > 4) and from PAT1 to PAT2 (Nm > 14) (Table 4).

**Table 4.** Effective numbers of migrants per generation (Nm) among *Spirochaeta* populations of the OTU6, OTU7 and OTU40.

| OTU | From | $\theta \pm se$ | To | $N_m \pm se$ |
| --- | --- | --- | --- | --- |
| OTU6 | KGI | $0.003 \pm 0.0003$ | KER | **$9.81 \pm 3.14$** |
| | | | PAT1 | $0.12 \pm 0.06$ |
| | | | PAT2 | $0.14 \pm 0.09$ |
| | KER | $0.004 \pm 0.0004$ | KGI | **$2.63 \pm 1.19$** |
| | | | PAT1 | $0.12 \pm 0.05$ |
| | | | PAT2 | $0.07 \pm 0.05$ |
| | PAT1 | $0.003 \pm 0.0004$ | KGI | $0.14 \pm 0.05$ |
|  |  |  | KER | **$0.43 \pm 0.15$** |
|  |  |  | PAT2 | **$24.99 \pm 7.10$** |
| | PAT2 | $0.005 \pm 0.0005$ | KGI | $0.20 \pm 0.07$ |
|  |  |  | KER | **$0.27 \pm 0.16$** |
|  |  |  | PAT1 | **$14.57 \pm 4.93$** |
| OTU7 | KGI | $0.002 \pm 0.0003$ | KER | **$1.39 \pm 0.60$** |
|  |  |  | PAT1 | **$0.64 \pm 0.23$** |
| | | | PAT2 | $0.03 \pm 0.02$ |
| | KER | $0.006 \pm 0.0006$ | KGI | **$0.42 \pm 0.13$** |
|  |  |  | PAT1 | **$24.41 \pm 8.23$** |
|  |  |  | PAT2 | **$2.00 \pm 0.54$** |
| | PAT1 | $0.015 \pm 0.0029$ | KGI | $0.16 \pm 0.06$ |
|  |  |  | KER | **$0.41 \pm 0.16$** |
|  |  |  | PAT2 | **$4.69 \pm 1.82$** |
| | PAT2 | $0.004 \pm 0.0005$ | KGI | $0.05 \pm 0.03$ |
| | | | KER | $0.13 \pm 0.09$ |
|  |  |  | PAT1 | **$19.95 \pm 9.57$** |
| OTU40 | KGI | $0.002 \pm 0.0002$ | KER | $0.09 \pm 0.04$ |
| | | | PAT1 | $0.01 \pm 0.01$ |
| | | | PAT2 | $0.02 \pm 0.02$ |
| | KER | $0.001 \pm 0.0002$ | KGI | **$0.87 \pm 0.21$** |
| | | | PAT1 | $0.02 \pm 0.02$ |
| | | | PAT2 | $0.02 \pm 0.02$ |
| | PAT1 | $0.005 \pm 0.0008$ | KGI | $0.01 \pm 0.01$ |
| | | | KER | $0.03 \pm 0.02$ |
|  |  |  | PAT2 | **$28.65 \pm 10.15$** |
| | PAT2 | $0.005 \pm 0.0008$ | KGI | $0.00 \pm 0.00$ |
| | | | KER | $0.06 \pm 0.02$ |
|  |  |  | PAT1 | **$15.94 \pm 10.48$** |

Only gene flows with $N_m$ values $> 0.25$ are considered as significant. Mean and standard error values were calculated from the 10 runs performed for each OTU.

The connectivity between the Patagonian and maritime Antarctic localities showed relatively low values of Nm, ranging from 0.001 (OTU40, from PAT2 to KGI) to 0.6 (OTU7, from KGI to PAT2) (Table 4). The OTU6 and OTU7 were both characterized by a substantial gene flow between maritime Antarctica and Kerguelen Islands that was stronger in the direction KGI to KER (OTU6, Nm ∼ 9.8 and OTU7, Nm ∼ 1.4) than in the direction KER to KGI (OTU6, Nm ∼ 2.6 and OTU7, Nm ∼ 0.4) (Table 4). Contrarily, an unidirectional and low gene flow from KER to KGI (Nm ∼ 0.9) was recorded for the OTU40 (Table 4). Finally, the connectivity between Patagonian (PAT1 and PAT2) and Kerguelen Islands (KER) localities was illustrated by three distinct patterns; a low-intensity flow (Nm < 0.5) predominant in the direction PAT1/PAT2 to KER for the OTU6, a substantial flow (Nm > 2) predominant in the direction KER to PAT1/PAT2 for the OTU7, and an absence of connectivity (Nm < 0.03) in the case of the OTU40 (Table 4).

### Contribution of contemporary selection versus historical processes in shaping the *Spirochaeta* microdiversity

For each of the three selected *Spirochaeta* OTUs, neutral ecological processes were essential in shaping the population composition turnover in *Abatus* gut membrane. According to the quantitative parsing of ecological processes, the composition of *Spirochaeta* population was mostly driven by ecological drift, ranging from 50% (OTU40) to 74% (OTU6) of turnover, followed by dispersal limitation, ranging from 12% (OTU6) to 20% (OTU40) of turnover, and homogenizing selection, ranging from 9% (OTU6) to 19% (OTU40) of turnover (Table 5).

**Table 5.** Quantitative parsing of ecological processes driving populations turnover within *Spirochaeta* OTUs.

| *Spirochaeta* OTU | Ecological processes contributions |  |  |  |  |
| --- | --- | --- | --- | --- | --- |
|  | Homogeneous selection (%) | Homogenizing dispersal (%) | Ecological drift (%) | Dispersal limitation (%) | Variable selection (%) |
| OTU6 | 2.7 | 8.8 | 74.0 | 12.1 | 2.3 |
| OTU7 | 0.4 | 10.3 | 63.7 | 17.6 | 8.0 |
| OTU40 | 0.3 | 19.4 | 49.6 | 21.7 | 9.1 |

According to the Stegen et al. (2013) approach, percentage refers to the percentage of pairs of communities that appear to be driven by either homogeneous selection, homogenizing dispersal, ecological drift, dispersal limitation or variable selection.

Overall, deterministic processes (*i.e.* homogeneous and variable selection) did not account for more than 10% of the populations’ turnover.

The MMRR approach was used to disentangle the relative effect of geographic distance environmental abiotic differences on the *Spirochaeta* oligotype compositions between samples. The geographic distance matrix was linearly correlated to the abundance-based similarity matrix of *Spirochaeta* population composition for OTU7 and OTU40, explaining about 31% and 67% of the observed variation, respectively (Table 6).

**Table 6.**
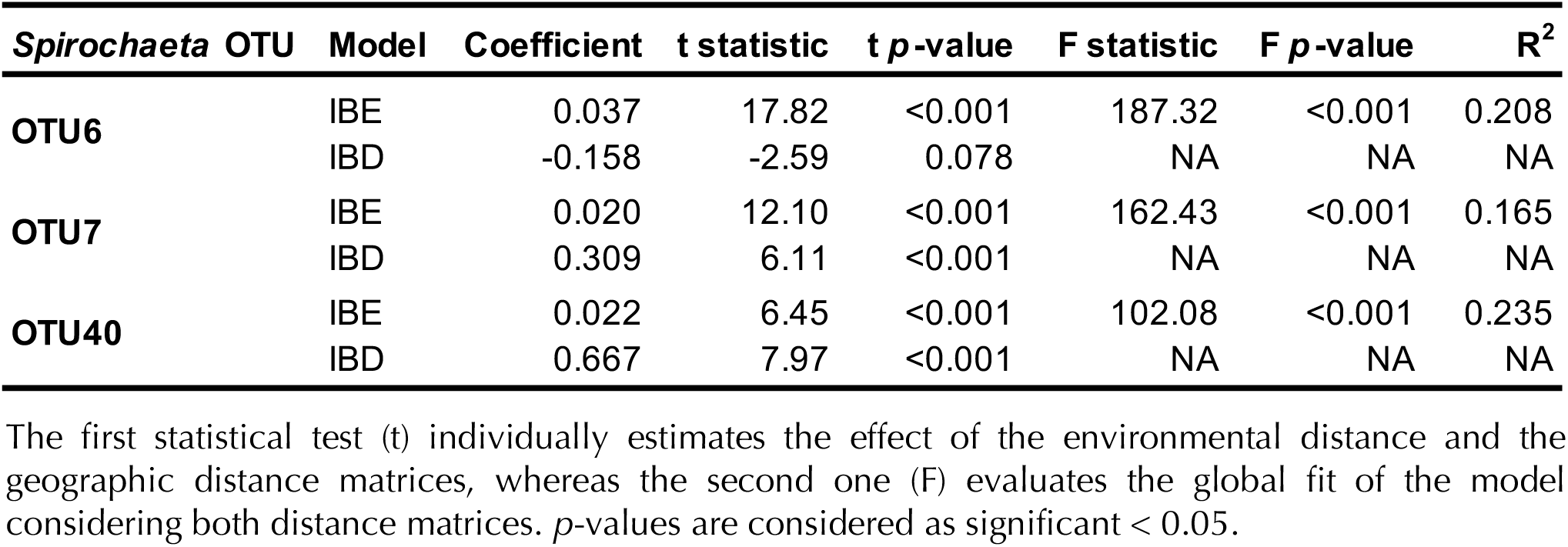
Multiple Matrix Regression with Randomization (MMRR) to quantify the relative effects of isolation by distance (IBD) and isolation by environment (IBE) on oligotypes assemblage within *Spirochaeta* OTUs.

| *Spirochaeta* OTU | Model | Coefficient | t statistic | t p-value | F statistic | F p-value | R<sup>2</sup> |
| --- | --- | --- | --- | --- | --- | --- | --- |
| OTU6 | IBE | 0.037 | 17.82 | <0.001 | 187.32 | <0.001 | 0.208 |
|  | IBD | -0.158 | -2.59 | 0.078 | NA | NA | NA |
| OTU7 | IBE | 0.020 | 12.10 | <0.001 | 162.43 | <0.001 | 0.165 |
|  | IBD | 0.309 | 6.11 | <0.001 | NA | NA | NA |
| OTU40 | IBE | 0.022 | 6.45 | <0.001 | 102.08 | <0.001 | 0.235 |
|  | IBD | 0.667 | 7.97 | <0.001 | NA | NA | NA |

The first statistical test (t) individually estimates the effect of the environmental distance and the geographic distance matrices, whereas the second one (F) evaluates the global fit of the model considering both distance matrices. *p*-values are considered as significant < 0.05.

In contrast, the geographic distance did not significantly impact *Spirochaeta* oligotype composition for OTU6 (Table 6). Whatever the OTU considered, the environment distance had a significant but slight contribution (< 4% of the observed variation) to the *Spirochaeta* population composition (Table 6), and the global R^2^ of the model (environmental and geographic distance) did not exceed 24%.

## Discussion

In this study, we coupled 16S rRNA metabarcoding and oligotyping algorithm to reveal the microdiversity within three bacterial OTUs affiliated to the *Spirochaeta* genus, and co-distributed across Patagonia, Kerguelen Islands, and maritime Antarctic provinces of the Southern Ocean. Through this innovative approach, we identified numerous oligotypes within each of the *Spirochaeta* OTUs. These oligotypes, corresponding to *Spirochaeta* sub-taxa, were characterized by contrasting geographic distribution and high levels of 16S rRNA gene similarity (> 97%). Taking advantage of the populational taxonomic resolution provided by the oligotype definition, we depicted the *Spirochaeta* biogeographic patterns across the analyzed provinces in the SO, using various tools adapted from population genetics classically applied in phylogeographic study of macroorganisms’ models. Despite its low substitution rate (approximately 1% in 50 million years [85]), our study demonstrates that the 16s rRNA gene still has its advantage in effectiveness and efficiency since it offers the best compromise between an informative genetic signal, and robust screening of global microbial diversity at intra-OTU level, in a wide range of barely unknown habitats [17].

Unlike the studies with macroorganisms, which are usually more demanding in terms of individual sampling effort, we benefit here from the high sequencing depth provided by the metabarcoding of a low-diversity habitat (*i.e.* the *Abatus* gut tissue). This allows a robust coverage of the *Spirochaeta* diversity (up to 180,000 sequences per OTU), and thus high precision of the oligotypes frequencies. Our methodology echoes the “metaphylogeographic” approach recently proposed by Turon *et al*. [86] to investigate eukaryotic intraspecies diversity through COI (cytochrome c oxidase subunit I) gene amplicon-sequencing and an oligotyping-like cleaning protocol of the reads based on entropy variation. We propose to expand this concept of “metaphylogeography” to the prokaryotes since our method permits phylogeographic inferences of uncultured microbes from a wide range of habitats.

The β-diversity analysis performed at the *Spirochaeta* genus level revealed that each of the three geographic provinces might host specific *Spirochaeta* OTUs representing distinct phylogenetic lineages. We also reported a non-random distribution trend with contrasting patterns of *Spirochaeta* OTU abundances across the localities. Nevertheless, about half of the *Spirochaeta* OTUs exhibited a broad distribution encompassing Patagonia, maritime Antarctica, and the Kerguelen Islands located more than 7,000 kilometers to the east. This result suggests that despite being mostly detected in *Abatus* gut, and to a lesser extent in marine benthic sediments, some *Spirochaeta* representatives would disperse through the SO currents. Concordantly, previous campaigns of high-throughput sequencing of the ocean water column have consistently reported the presence of free-living *Spirochaeta* OTUs in surface to mesopelagic water, away from the coastlines [87].

For each of the three assessed OTUs, the *Spirochaeta* populations were expected to be remarkably homogeneous between the two Patagonian sites due to the geographic vicinity and the absence of an evident oceanographic barrier. Indeed, most of the *Spirochaeta* oligotypes were shared between these two localities that displayed the lowest genetic and phylogeographic structure for each of the three OTUs. High levels of gene flow, illustrated by a high effective number of migrants per generation (Nm), were also recorded between the two Patagonian localities, in accordance with the homogeneity of their oligotype compositions. Similarly, low or absent differentiation patterns along the Atlantic coast of Patagonia were previously reported for marine Patagonian macroorganisms, including notothenioid fishes [88] scorched mussels [89], and pulmonate gastropods [90]presumably due to their high dispersal potential and the ecological continuum of the sampled localities that may conform a same biogeographic province connected through the equator-ward Falkland current [91, 92]. Further phylogeographic studies focusing on microbial taxa of additional sampling sites from Atlantic Patagonia should confirm the microbial biogeographic consistency of this province.

Between Patagonian and maritime Antarctic provinces, *Spirochaeta* populations exhibited strong genetic and phylogeographic structure, as illustrated by the high F_ST_ and ɸ_ST_ values and the relatively low number of shared oligotypes. Additionally, low levels of gene flow were estimated between these two provinces (*i.e.* effective numbers of migrants per generation Nm < 1). These results corroborate our hypothesis that the APF hinders individual dispersion and genetic homogeneity among bacterial populations and suggest that the geographically structured *Spirochaeta* populations from these two provinces are genetically diverging over time [93, 94]. Previous studies focusing on diverse macroorganisms taxonomic groups of the SO have evidenced the critical role of the APF on biogeographic patterns, as an open-ocean barrier inducing a genetic break between South America and Antarctica (*e.g.* ribbon worms, [95]; brittle stars, [96]; notothenioid fishes, [25, 97]; limpets [26, 29, 98]; sea urchins [24]). Regarding the microbial distribution patterns, significant β-diversity differences between prokaryotes assembly from both sides of the APF have been reported in the past, but most of the studies focused on global community in the water column, at high taxonomic resolution (summarized in Flaviani et al. [20]). Here, we extend this discontinuity in bacterial diversity to a finer taxonomic resolution (*i.e.* intra-OTU), revealing province-restricted oligotypes and strong genetic and phylogeographic structure between Patagonian and maritime Antarctic *Spirochaeta* populations.

Contrarily, and despite the substantial geographic distance separating the sub-Antarctic Kerguelen Islands and the Patagonian and Antarctic sites (> 6,500 kilometers), population genetic analyses suggest the existence of some level of connectivity between Kerguelen and the other sites. These findings support a potential dispersion of *Spirochaeta* taxa from Patagonia and maritime Antarctica to the Kerguelen Islands, and contrariwise, from the Kerguelen Islands to Patagonia and maritime Antarctica. As evidenced by the numerous shared oligotypes, such connectivity would maintain a sufficient gene flow among these provinces to partially counteract the genetic divergence driven by selection, mutation, and genetic drift, inducing oligotypes mixing, and limiting the spatial differentiation of oligotypes assembly [1] [99]. Moreover, we suggest that this gene flow is not bidirectional, but governed by exclusively eastward oriented dispersion routes (Figure 5), following the major and constant flow of the ACC [100]. Under this scenario, *Spirochaeta* individuals from Kerguelen Islands may seed towards Patagonia following the ACC eastward flow around Antarctica. Such ACC-mediated connectivity among sub-Antarctic provinces (Patagonia and Kerguelen Islands) is well known in a wide range of benthic macroorganisms populations, such as buoyant kelps *Durvillaea antarctica* [38] and *Macrocystis pyrifera* [101], and several kelp-associated macroinvertebrates [32, 34, 35, 100, 102].

**Figure 5.**
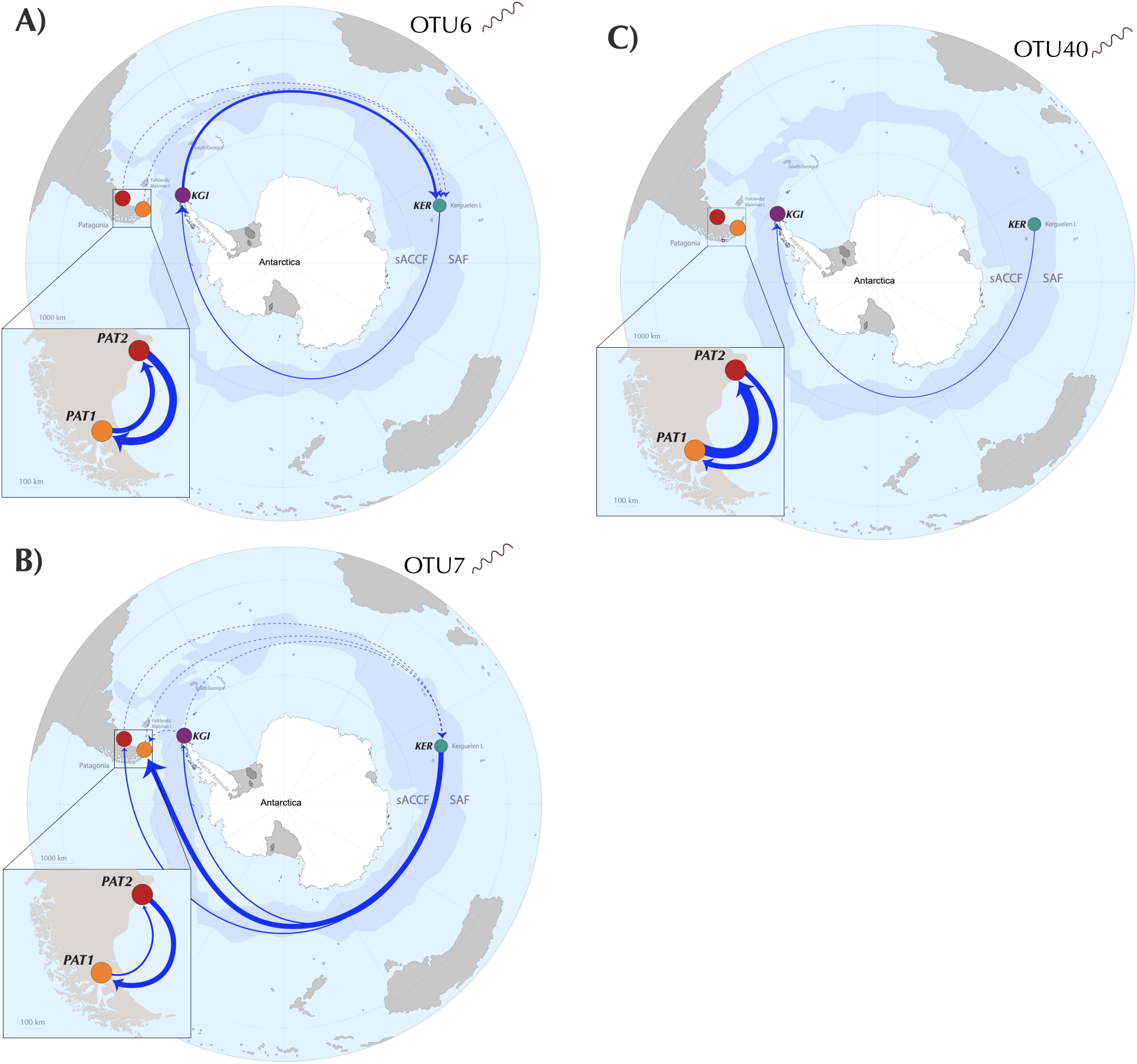
Gene flows summary and proposed dispersal routes across the Southern Ocean for each *Spirochaeta* taxa. Only the gene flows with Nm values > 0.25 are graphically represented. Discontinuous arrows represent Nm values > 0.25 and < 1, continuous arrow represent Nm values > 1. Continuous arrows’ width is proportional to Nm values.

Occasionally, *Spirochaeta* individuals from Kerguelen Islands may also be able to reach the maritime Antarctic province. Such pattern has been recently reported for the southern bull kelp *D. antarctica*, a typical sub-Antarctic macroalgae, which is transported by rafting to as far as the West Antarctic Peninsula coasts, pushed by the circumpolar flow of the ACC or by storms leading to the occasional crossing of the APF [103].

Several studies have provided evidence of a high dispersal capacity of marine bacteria by comparing community composition mostly at high taxonomic resolution (*e.g.* class, genus, or OTU) among various water masses and oceanic regions [104–107]. Particularly, the most abundant marine bacteria are supposed to migrate between adjacent regions through passive transport [104]. However, this is the first time that it is evidenced in bacterial populations through the microdiversity resolution, being solely suggested so far at the community level [20]. An innovative conceptual framework called “Microbial Conveyor Belt” (MCB) has been proposed by Mestre & Höfer [108], to emphasize that the marine microorganisms’ dispersion would not merely rely on passive and stochastic dispersal, but instead on the adaptation of life-history traits (*e.g.* dormancy stage). These traits would allow microorganisms to successfully and recurrently disperse in unfavorable habitats through specific dispersion avenues [109]. Here, we provided empiric results from *Spirochaeta* population based on genetic data supporting a partial MCB in the SO driven by the ACC. Unfortunately, details about the benthic *Spirochaeta* taxa’s ecology are scarce, with a single isolated strain from subseafloor sediment [110]. Thus, the life-history traits of *Spirochaeta*, as the sporulation capacity, remain to be investigated to further understand its distribution pattern in the SO. Nevertheless, in order to disperse, we propose that *Spirochaeta* individuals (enriched in the digestive tract) could be released from the host gut towards the surrounding benthic sediments through fecal pellets. Such enrichment of the digesta with taxa from the host microbiota, as well as the presence of *Spirochaeta* within the fecal material, have been demonstrated in the sea urchin species *Lytechinus variegatus* [111]. The released *Spirochaeta* individuals may be resuspended in the water column through the action of one or several processes such as upwelling, bioturbation by the benthic deposit-feeders, or water column mixing during winter [108, 112, 113]. Once in the water column, these *Spirochaeta* individuals may disperse over large geographic scales, transported through oceanographic features (*e.g.* currents, punctual meteorological events) [108]. The attachment to suspended particulate matter, either biotic (*e.g.* hitchhiking on zooplankton [114] and seaweed [115]) or abiotic (*e.g.* microplastics [116], known to have a long-distance dispersion potential), may also contribute to the bacterial spreading in the oceans [107, 108].

The marine prokaryote communities are usually considered widely dispersed and mainly shaped by contemporary ecological processes such as environmental filtering [106, 117]. By applying the ecological framework developed by Stegen et al. [71] to oligotypes data, we found contrarily that ecological drift was the predominant stochastic mechanism shaping intra-populations turnover within *Spirochaeta* taxa across the SO. Our previous study of the *Abatus* gut microbiota showed that non-neutral processes drove the bacterial community at the OTU level in the host gut tissue [54]. While deterministic processes are usually prevalent in structuring microbial communities’ assembly at a higher taxonomic resolution [18, 79, 118, 119], the stochastic mechanisms tend to have a more significant contribution at finer taxonomic scales [105], since niche overlapping and functional redundancy enhance the susceptibility of populations to drift [120]. Thus, the biogeographic structure observed within *Spirochaeta* populations might result from stochastic birth, death, disturbance, emigration, and immigration events rather than oligotype-sorting through the biotic and abiotic environmental variations [1, 121, 122]. Consistently, the MMRR analysis revealed that the Isolation-by-Environment (IBE) model might account for a low percentage of the *Spirochaeta* oligotypes turnover. Altogether, these results validate the strategy applied in our study, that is, to focus on specialist bacterial taxa hosted in sibling sea urchin species with the same habitat preferences, in order to homogenize the environment, to reduce the diversity, to soften the deterministic selection driven by environmental variations and to maximize the detection of neutral micro-evolutive processes associated with biogeography [123].

By analogy with the genetic drift, whereby changes in gene frequencies occur solely by chance in a population [76], our result suggests that the microdiversity observed within the *Spirochaeta* taxa would be mostly generated by genetic drift without any adaptive implications. An earlier study reported that microdiversity observed in the 16S rRNA gene of marine coastal *Vibrio splendidus* isolates was ecologically neutral [124]. Nevertheless, we cannot discard that, while the microdiversity within the V4-V5 of 16S rRNA gene-targeted here is likely to be acquired through neutral processes [125], it may also be associated with substantial modifications in niche-defining traits and functional attributes specific of the *Spirochaeta* strains, driven by deterministic processes, in order to cope with local conditions [17, 126]. Further studies will need to focus on other loci (*e.g.* functional genes), potentially under selection, as they are expected to display a higher degree of differentiation among populations and to provide an insight into the ecology of the *Spirochaeta* sub-taxa [127]. Furthermore, notwithstanding the consistency of the global phylogeographic and connectivity patterns depicted across the three tested OTUs, we also reported some differences according to the taxa considered, which might be related to different ecotypes with distinct ecological niches or different dispersal capacity. For instance, various marine bacterial taxa, such as the cyanobacteria *Synechococcus* or the *Vibrio* populations, demonstrate fine-tuning of their physiology by accumulating microdiversity in functional genes through duplication events, SNPs, and allelic variants [128, 129]. Alternatively, these differences may also be related to the *Spirochaeta* OTU abundance, since the more relatively abundant the *Spirochaeta* populations were (*i.e.* higher number of sequences retrieved from the gut tissue through the metabarcoding approach), the more they tend to exhibit cosmopolitan oligotypes (*i.e.* detected across each of the four localities). It is not unreasonable to infer that a larger population may have more chance to migrate and successfully reach a suitable habitat, while small-size populations may be more likely diluted along the dispersal route with no/too few dispersive particles to establish in the new habitat [104].

The diversity units defined by 16S rRNA gene sequences are generally considered as insensitive to diversification resulting from dispersal limitation [10]. Contrastingly, we reported that the dispersal limitation was the second most crucial ecological factor driving the turnover of *Spirochaeta* oligotypes, and by extension, their genetic divergence. Dispersal limitation is classically considered as a historical factor since current oligotypes assemblage results from past dispersal limitations [1]. Our result indicates that the potentially suitable habitats are too distant [130], or inaccessible due to the existence of oceanic currents [104, 131], hence limiting the homogenization of *Spirochaeta* oligotypes’ frequencies across populations and allowing the neutral genetic divergence of genomic regions overtime via genetic drift [10, 132]. Note that our results obtained from distinct methodologies (*i.e.* the genetic differentiation and phylogeographic structure, the contribution of dispersal limitation from Stegen *et al.* framework [71], and the contribution of the geographic distance (IBD) from the MMRR analysis) were highly consistent with each other, and across the three selected *Spirochaeta* OTUs. For instance, the OTU40 that harbored the overall highest value of genetic divergence was also characterized by the highest estimated contribution of geographic distance and dispersal limitation, thus supporting the interrelation between genetic divergence and oligotypes population turnover, and the overall consistency of the approach implemented.

## Conclusion

Our study highlights the application of V4-V5 16S rRNA gene metabarcoding and oligotyping approach as rapid, robust, and resolutive enough to unravel marine bacterial phylogeographic patterns and detect genetic connectivity among the SO provinces. Taken together, the three *Spirochaeta* OTUs analyzed evidence three consistent phylogeographic patterns, classically observed in the studies involving benthic macroinvertebrates across the SO: (i) a high populational and genetic homogeneity within the Patagonia province, (ii) a strong barrier to dispersal between Patagonia and maritime Antarctica due to the APF, resulting in a high differentiation of *Spirochaeta* populations, and (iii) the existence of connectivity between sub-Antarctic provinces of the Kerguelen Islands and Patagonia, and from Kerguelen Islands to maritime Antarctic, due to the ACC-mediated connectivity. Nevertheless, as connected as these provinces are, the gene flow does not seem to be strong enough to prevent the ongoing intraspecific differentiation process of the *Spirochaeta* taxa. The microdiversity of *Spirochaeta*, underlying these biogeographic patterns, is essentially driven by historical processes, such as ecological and genetic drift, and dispersal limitation related to the SO’s oceanographic features. In the future, extending this framework to other localities and taxonomic groups will contribute to the comprehensive understanding of the Southern Ocean microbiota.

## Supporting information

Additionnal 1

Additionnal 2

Additionnal 3

Additionnal 4

Additionnal 5

## Declaration

### Ethics approval and consent to participate

Not applicable

### Consent for publication

Not applicable

### Availability of data and materials

The datasets generated for this study can be found at the National Centre for Biotechnology Information (NCBI) repository, under the following accession numbers: PRJNA658980, PRJNA590493 and PRJNA659050, corresponding to the datasets of *Abatus cordatus* from Kerguelen Archipelago, *Abatus agassizii* from West Antarctic Peninsula and *Abatus cavernosus* from South America, respectively.

### Authors’ contributions

EP, JO and LC designed the study. EP and JO organized the sampling missions. EP, JO, LC and GS collected samples. GS extracted the DNA, organized sequencing and managed data mining and analyses. NS contributed to the gene-flow and the MMRR analyses and designed the illustrative maps. GS, JO, NS, CGW and EP interpreted the results. GS wrote the manuscript. All authors contributed substantially to manuscript revisions. All authors read and approved the final manuscript.

## Acknowledgements

We thank Dr Peter Beerli and Dr Franz-Sebastian Krah for their technical supports. We also thank Jonathan Flores for providing *Abatus* samples from Argentinian Patagonia, and Dr Karin Gérard for managing the sampling logistic in Chilean Patagonia. We recognize the Chilean Antarctic Institute (INACH) for the logistic support during the Chilean Antarctic Expeditions (ECA 55 and 56).

## Supplementary material

**Additional file 1.** Pairwise PERMANOVA on *Spirochaeta* OTUs composition dissimilarities among localities. *p*-values are adjusted using the default Bonferroni method implemented in the *pairwiseAdonis* R package and are considered as significant < 0.05.

**Additional file 2.** Abundance and closest sequence retrieved from Blast analysis for each of the three OTUs analysed through the MED pipeline.

(1) Bowman JP, McCuaig RD: Biodiversity, community structural shifts, and biogeography of prokaryotes within Antarctic continental shelf sediment. *Appl Environ Microbiol* 2003, 69:2463-2483.

(2) Acosta-González A, Rosselló-Móra R, Marqués S: Characterization of the anaerobic microbial community in oil-polluted subtidal sediments: aromatic biodegradation potential after the Prestige oil spill. *Environmental microbiology* 2013, 15:77-92.

**Additional file 3.** Relative contribution of each locality in the total abundance of OTU6, OTU7 and OTU40 sequences. Colors are assigned to the different localities.

**Additional file 4.** Summary of number of oligotypes in each locality and per OTU of *Spirochaeta*.

**Additional file 5.** Accumulation curves of OTU6 (A), OTU7 (B) and OTU40 (C) oligotypes richness. Colors are assigned to each locality. Extrapolation is calculated from Hill numbers of richness (q=0).

