## Supplementary material for "Exploring the microdiversity within marine bacterial taxa: Towards an integrated biogeography in the Southern Ocean": Additionnal 1

### Slide 1
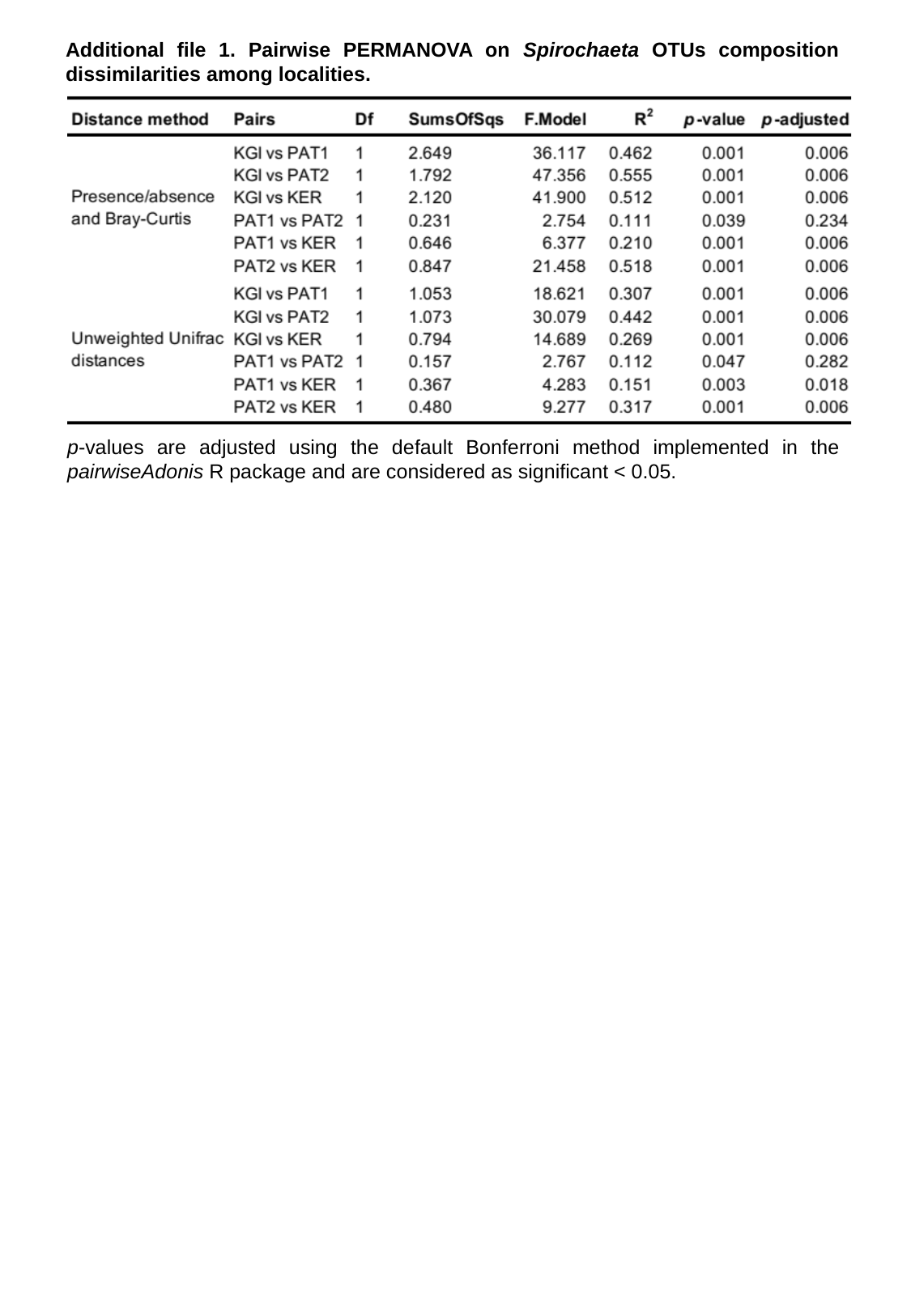

Additional file 1. Pairwise PERMANOVA on Spirochaeta OTUs composition dissimilarities among localities.
p-values are adjusted using the default Bonferroni method implemented in the pairwiseAdonis R package and are considered as significant < 0.05.
