## Supplementary material for "Exploring the microdiversity within marine bacterial taxa: Towards an integrated biogeography in the Southern Ocean": Additionnal 2

### Slide 1
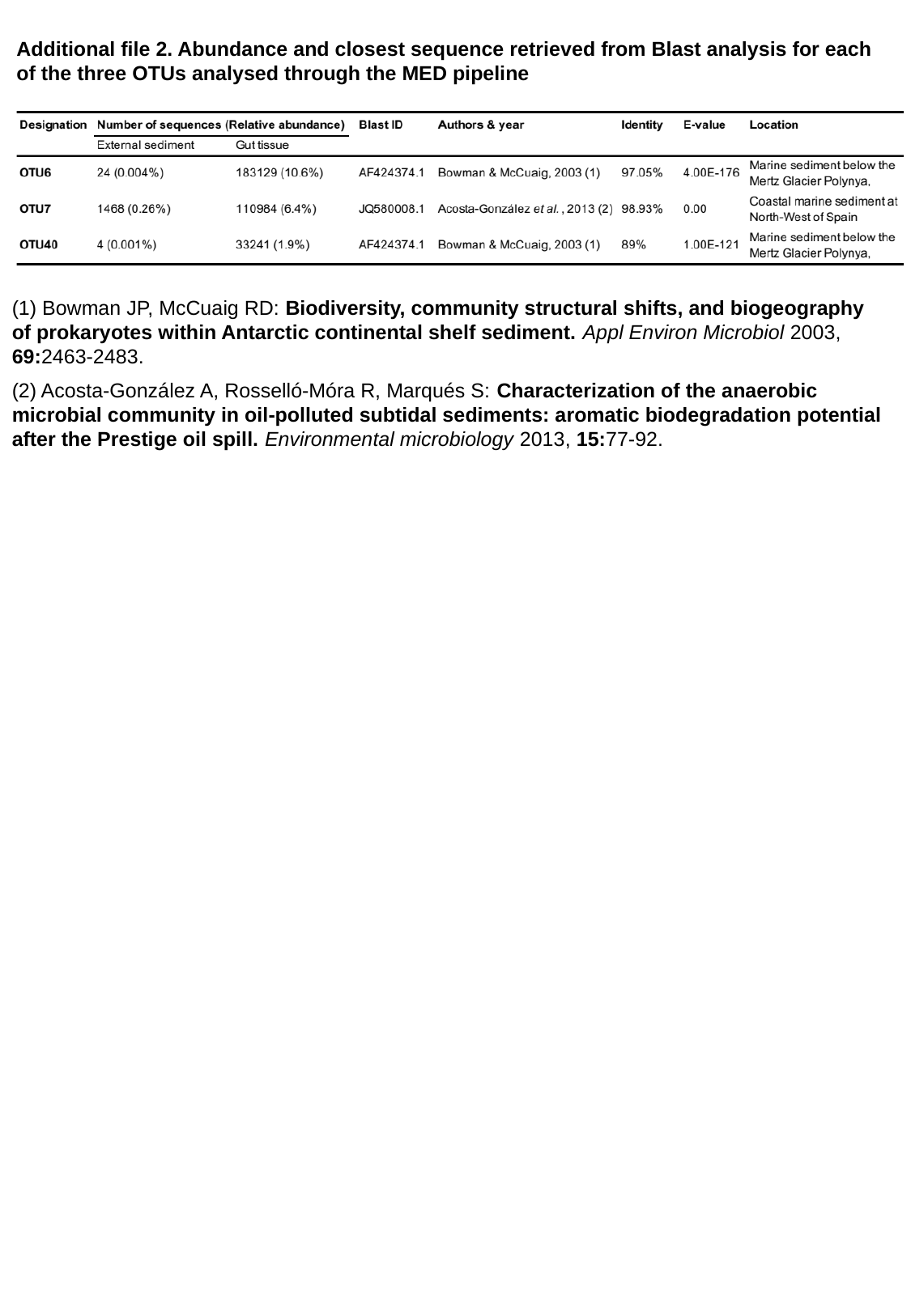

Additional file 2. Abundance and closest sequence retrieved from Blast analysis for each of the three OTUs analysed through the MED pipeline
(1) Bowman JP, McCuaig RD: Biodiversity, community structural shifts, and biogeography of prokaryotes within Antarctic continental shelf sediment. Appl Environ Microbiol 2003, 69:2463-2483.
(2) Acosta‐González A, Rosselló‐Móra R, Marqués S: Characterization of the anaerobic microbial community in oil‐polluted subtidal sediments: aromatic biodegradation potential after the Prestige oil spill. Environmental microbiology 2013, 15:77-92.
