## Supplementary material for "Exploring the microdiversity within marine bacterial taxa: Towards an integrated biogeography in the Southern Ocean": Additionnal 3

### Slide 1
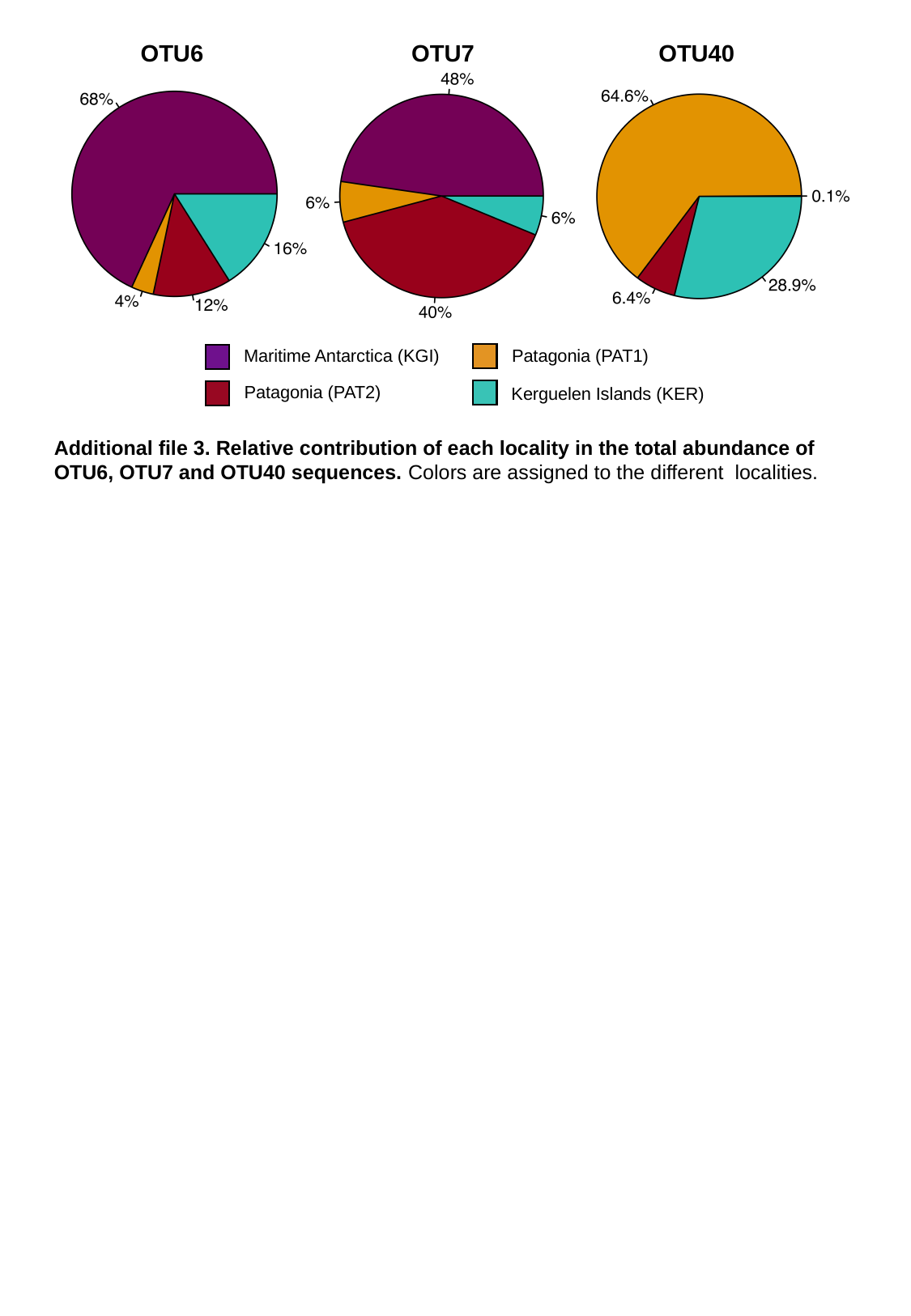

OTU6
OTU7
OTU40
Patagonia (PAT1)
Maritime Antarctica (KGI)
Patagonia (PAT2)
Kerguelen Islands (KER)
Additional file 3. Relative contribution of each locality in the total abundance of OTU6, OTU7 and OTU40 sequences. Colors are assigned to the different localities.
