## Supplementary material for "Exploring the microdiversity within marine bacterial taxa: Towards an integrated biogeography in the Southern Ocean": Additionnal 4

### Slide 1
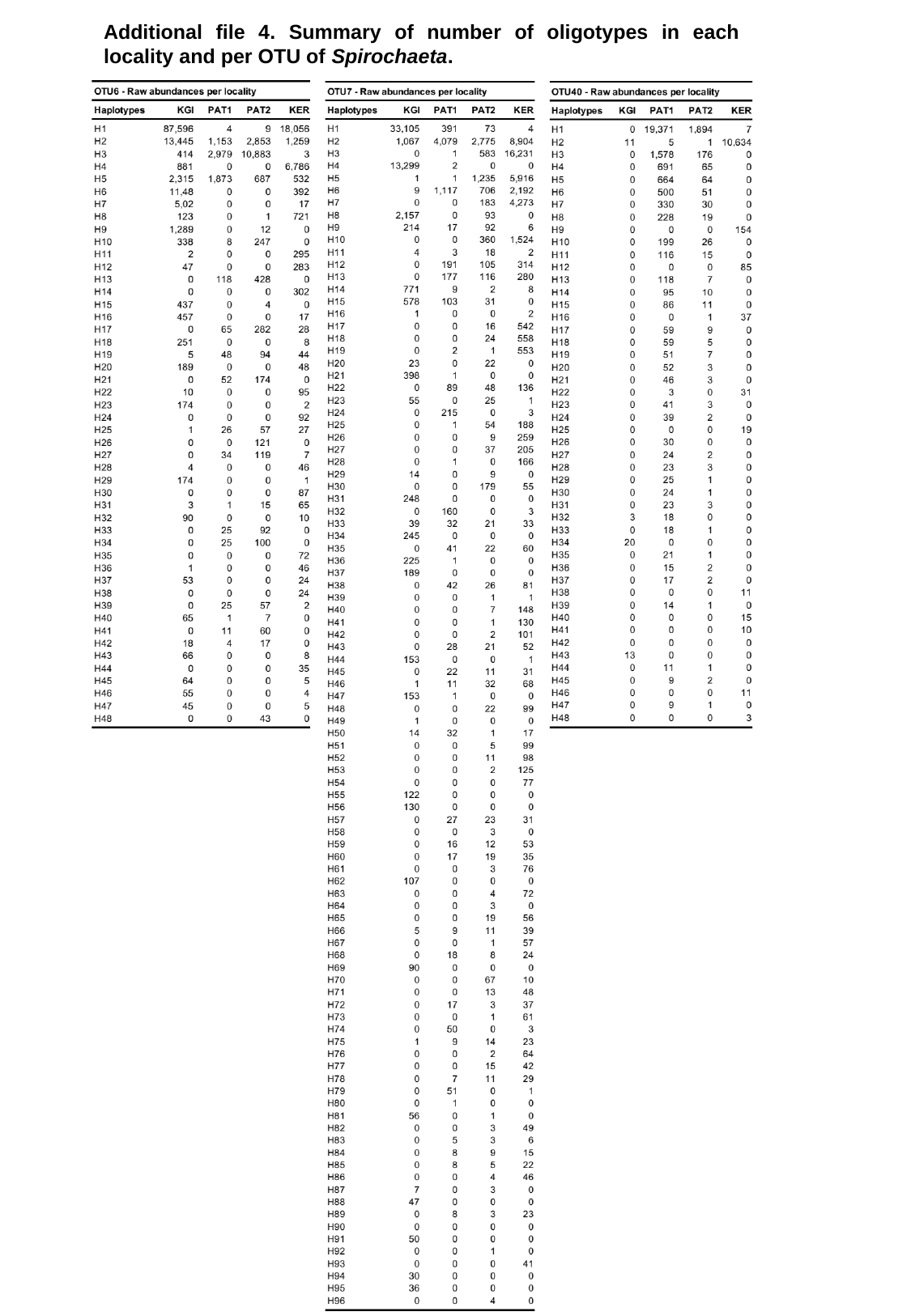

Additional file 4. Summary of number of oligotypes in each locality and per OTU of Spirochaeta.
