## Supplementary material for "Exploring the microdiversity within marine bacterial taxa: Towards an integrated biogeography in the Southern Ocean": Additionnal 5

### Slide 1
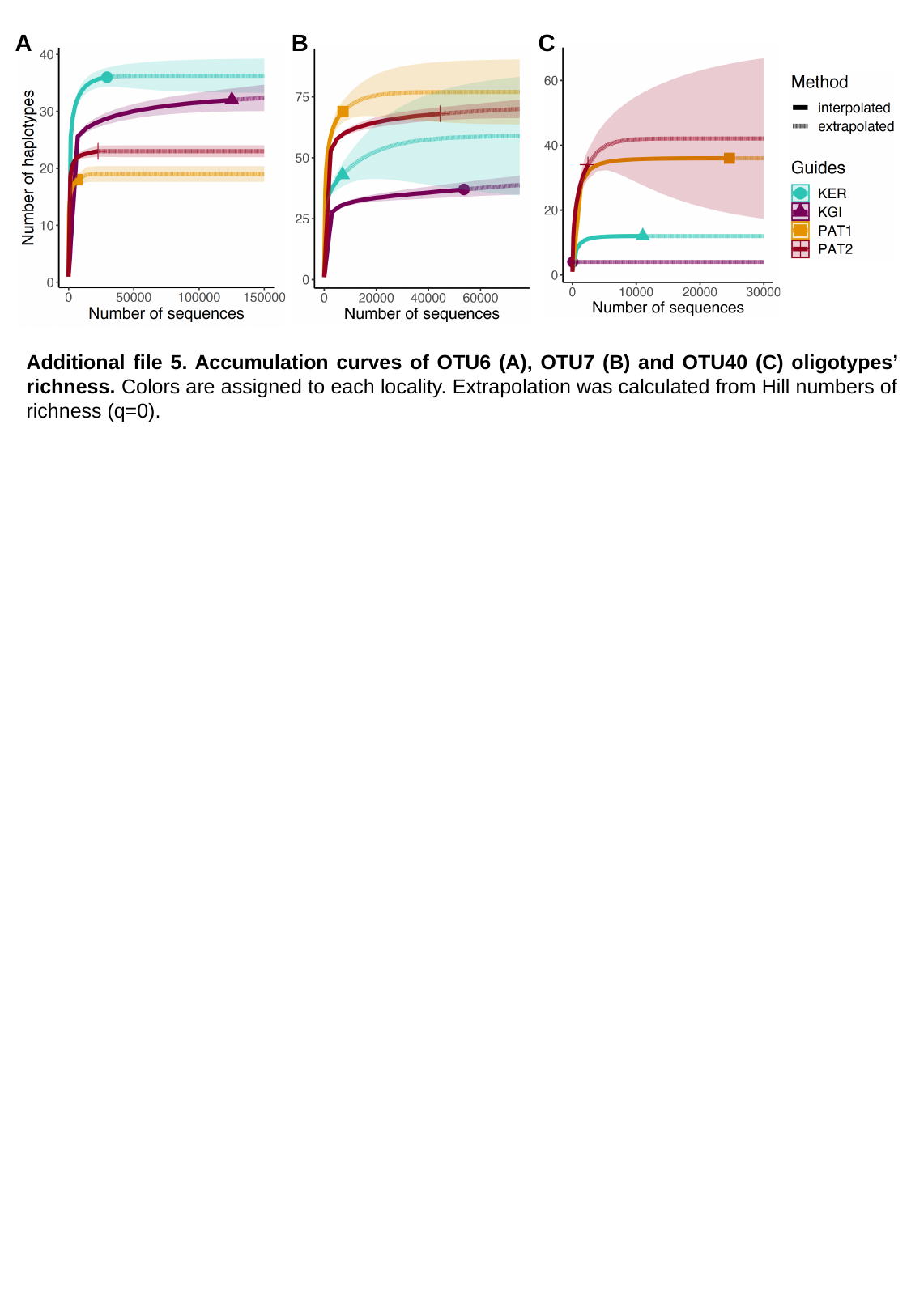

A
B
C
Additional file 5. Accumulation curves of OTU6 (A), OTU7 (B) and OTU40 (C) oligotypes’ richness. Colors are assigned to each locality. Extrapolation was calculated from Hill numbers of richness (q=0).
